# Spatial coding supports auditory conceptual navigation

**DOI:** 10.1101/2025.02.04.636440

**Authors:** Kyle Jasmin, Max Bullock, Frederic Dick, Roger Atkins, Roberto Bottini, Federica Sigismondi, Adam Tierney

**Affiliations:** Royal Holloway, University of London; University College London, University of London; University of Trento; Birkbeck, University of London

## Abstract

Grid cells in human entorhinal cortex encode spatial layouts for real-world navigation, yet their role in conceptual navigation remains unclear. Here we show that mentally transforming tones within a purely auditory pitch-duration space engages spatial circuits, and that such resources are causally necessary. In Experiment 1, participants trained for five days to navigate through a purely conceptual pitch-duration auditory space, then underwent fMRI on Day 6. We observed a six-fold modulation of entorhinal BOLD signals aligned to each participant’s trajectory angles, similar to grid-cell firing in physical space. Stronger grid-like coding predicted larger training-related gains. In Experiment 2, a new cohort performed the same task under either a spatial or non-spatial interference load. Only the spatial condition selectively disrupted performance on trials requiring mental “movement,” indicating a causal reliance on spatial resources. These findings provide evidence that auditory conceptual transformations recruit—and depend on—spatial grid-like computations in the entorhinal-hippocampal system, pointing to a domain-general role for spatial coding in organizing new knowledge along continuous dimensions.

## Introduction

Our survival depends crucially on the ability to mentally represent and navigate our environment, a capacity supported by the hippocampal-entorhinal system. Seminal rodent studies identified two principal cell types involved in spatial navigation: place cells—neurons in the hippocampus that fire in response to specific locations (O’Keefe & Dostrovsky, 1971)and grid cells in the entorhinal cortex, which exhibit a hexagonal firing pattern across a navigable space (Hafting et al., 2005). Grid cells have also been observed in humans using fMRI (Doeller et al., 2010; Horner et al., 2016); together with other cell types such as head direction cells (Muller et al., 1996), they allow us to encode “cognitive maps” of our environment (Behrens et al., 2018; Moser et al., 2008; Tolman, 1948).

Grid cell-like firing patterns (henceforth “grid-like coding”) appear not only in physical navigation but also in cognitive “movement” through abstract domains defined by continuous dimensions (Bellmund et al., 2018). For example, when participants navigated a conceptual “bird” space composed of visual features of cartoon birds (whose legs and neck varied in length, orthogonally), activity in entorhinal cortex displayed a six-fold symmetry characteristic of grid cells (Constantinescu et al., 2016). Similar findings have been demonstrated for olfactory space composed of two scents of varying concentrations (Bao et al., 2019; Raithel et al., 2023) social spaces (Park et al., 2021), and value spaces during prospective decision-making (Nitsch et al., 2024). These findings suggest that the hippocampal-entorhinal system may repurpose spatial coding mechanisms to support the organization and traversal of non-physical spaces. However, it remains unknown whether spatial resources are merely an epiphenomenon or whether they *causally support* conceptual navigation. Existing evidence comes mostly from correlational fMRI showing six-fold BOLD modulations in entorhinal cortex during movement through conceptual spaces. Testing whether spatial mechanisms are *necessary* for abstract navigation requires overloading or disrupting them while observing the consequences for task performance.

Here, we sought to test whether spatial navigation plays a causal role in conceptual reasoning by examining whether mentally “moving” through a purely auditory stimulus space is selectively impeded by a secondary spatial task. This space was defined by two auditory dimensions—pitch and duration—yielding a quasi-spatial framework that can be mentally traversed without relying on explicit spatial (e.g., higher/lower) language. There were several reasons for choosing an auditory space. First, there is already compelling evidence that auditory dimensions and spatial processing are closely linked in the human mind. For example, spatial linguistic metaphors for sound dimensions (a *high* pitch; a *long* note) predominate in many languages (Lakoff & Johnson, 2008), and psychophysical evidence demonstrates mental associations between space and auditory dimensions such as pitch (Dolscheid et al., 2013, 2014; Hartmann, 2017; Lidji et al., 2007), raising the question of whether cognitive resources are shared between auditory dimensions and space. Second, animal models already suggest entorhinal grid cells can code an auditory space, at least a one-dimensional one: single unit recordings in rats have shown that the same entorhinal neurons that exhibited grid-like coding during physical navigation were also involved with coding specific auditory frequencies, in a task where the animal needed to “navigate” to a specific frequency using a lever, to receive a reward (Aronov et al., 2017). Third, in human communication systems such as spoken language and music structural features are differentiated by multiple auditory dimensions simultaneously (Jasmin, Dick, Holt, et al., 2020; Jasmin, Dick, Stewart, et al., 2020; Jasmin et al., 2021, 2023). If integrating these dimensions, reasoning about them, or producing mental (auditory) imagery across them recruits space-like neural resources, this would require extension of current neurobiological models of music and language processing (Hagoort, 2019; Jasmin et al., 2019; Vuust et al., 2022). Thus, to test whether mental navigation in auditory space recruits domain-general spatial mechanisms, we conducted two complementary experiments.

In Experiment 1, we asked whether mental navigation within a purely auditory “pitch-duration” space (Figure 1) recruits neural coding mechanisms for spatial navigation. We trained 35 participants over five days to “move” through a two-dimensional pitch–duration space, then scanned them on Day 6 using fMRI. On each trial, participants heard an initial “reference tone”, followed by short spoken instructions—for example “pitch plus one, duration same”—directing them to a new tone they should imagine . They then heard a probe tone and indicated with a keypress whether it matched what they had imagined or not. We employed a standard grid-like coding analysis (Doeller et al., 2010) to test for six-fold modulation of BOLD responses in entorhinal cortex during the portion of each trial when participants actively imagined moving from one tone to another. We predicted that entorhinal activity would reflect a six-fold symmetry aligned to each participant’s movement angles in the auditory domain. We further explored whether individual differences in grid-like coding strength correlated with training-related improvements, self-reported task difficulty, and musical background (years of formal training and performance experience).

**Figure 1.**
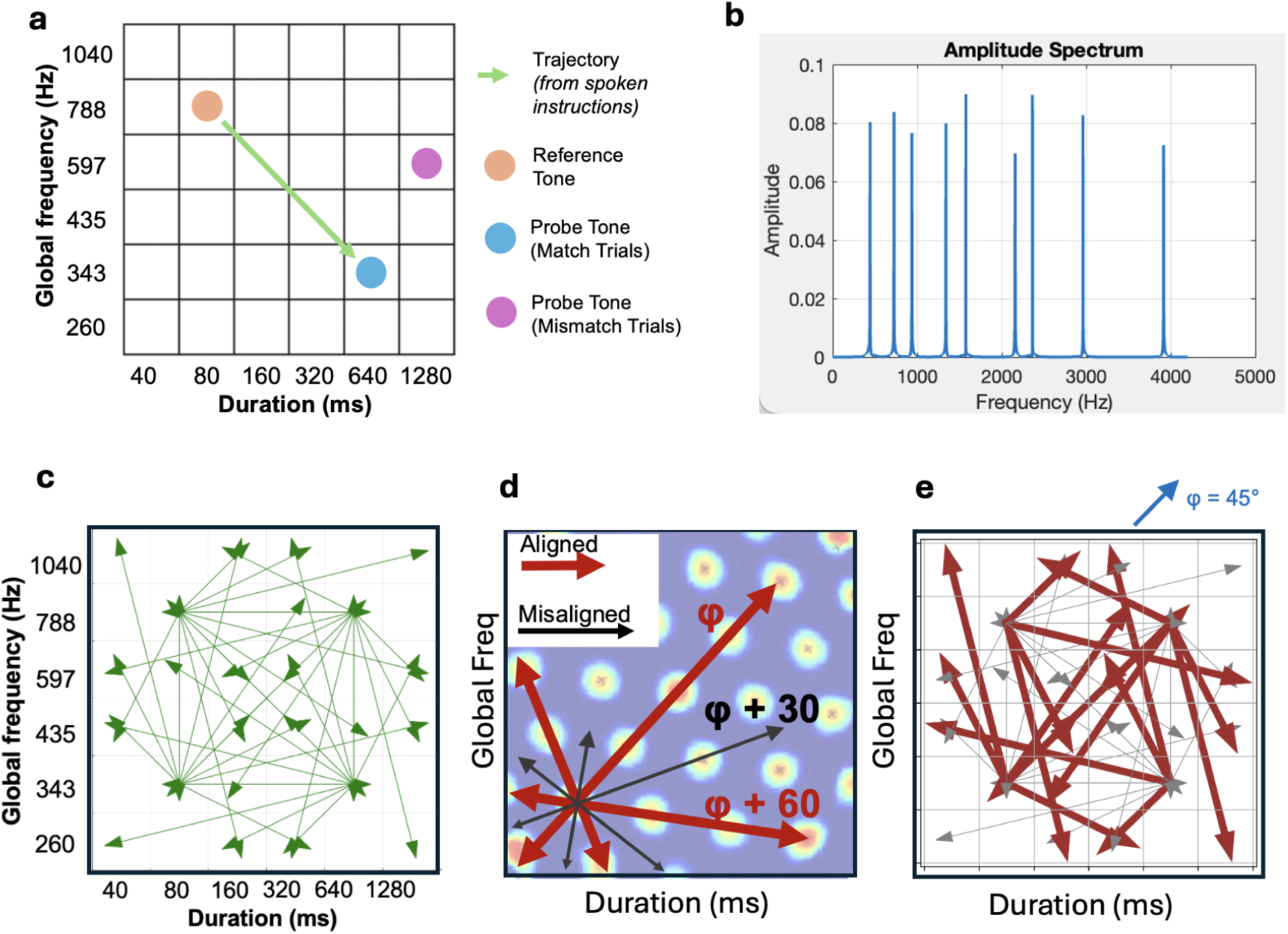
Stimuli and conceptual framework. **(a)** A 6×6 pitch–duration space, with an example trajectory (green arrow). The orange circle is the reference tone, the blue circle is a matching probe, and the pink circle is a mismatched probe. **(b)** Amplitude spectrum of an example tone (global frequency = 788 Hz). **(c)** The 48 stimulus trajectories. **(d)** Illustration of grid-like firing fields (background), showing example aligned trajectories (red) and a misaligned one (black), relative to grid angle φ. **(e)** Example of how trajectories in the space can be classified as Aligned (red) or Misaligned (grey) in the fMRI analysis, relative to an example grid orientation (φ) of 45°.

In Experiment 2, we examined whether spatial cognitive resources are causally necessary for this form of conceptual navigation. A new cohort performed the same auditory navigation task, but now under one of two concurrent interference conditions: a spatial load (judging if three letters are arranged left-to-right on a QWERTY keyboard) or a non-spatial load (judging if letters are in alphabetical order). We also manipulated whether traversal of the pitch-duration space was required (Move condition) or no (No-Move condition). Our causal inference relies on the assumption that checking the relative locations of keys on a QWERTY keyboard requires scanning 2D visual space, which has been shown to employ entorhinal grid-like coding (Julian et al., 2018; Nau et al., 2018; Sigismondi et al., 2024).

Alphabetical order is by contrast not routinely spatialized nor has it been shown to recruit the hippocampal-entorhinal system (Attout et al., 2014; Jonas et al., 2011). We predicted that imposing a spatial load would selectively disrupt performance on the auditory task only when participants had to “move” through the pitch–duration space—evidence for a domain-general reliance on spatial neurocognitive mechanisms. Full methodological details, including participant recruitment, stimuli, fMRI acquisition parameters, and analytical procedures, are provided in the Methods section.

## Results

### Brain Imaging of Left Entorhinal Cortex (Experiment 1)

As expected, participants’ accuracy improved over the 5-day at-home training (Figure 2a). We examined neural activity in the medial temporal lobe to identify any six-fold coding in entorhinal cortex, using a standard approach (Constantinescu et al., 2016; Doeller et al., 2010; Stangl et al., 2017). A quadrature filter localiser within a bilateral entorhinal cortex mask tested for the presence of six-fold symmetry voxel-wise without regard to grid angle (see GLM0 in Methods). This revealed two significant clusters, surviving a minimum extent threshold of 18 voxels (calculated using AFNI’s *3dClustSim* with ACF model, 10,000 iterations): one cluster in left entorhinal cortex (43 voxels) and one in right entorhinal cortex (33 voxels; Figure 2b). We analyzed the clusters separately. The preferred grid angle in left entorhinal cortex for each participant was estimated from runs 1 and 3 (see *GLM1* in Methods), weighted by the amplitude of the six-fold signal to ensure that the voxels with strongest signal contributed most to the grid angle estimation (Stangl et al., 2017). We then found that in independent runs 2 and 4 activity during trials where the trajectory was aligned with the grid angle estimated in runs 1 and 3 (i.e. ±15 degrees) was greater than during trials where the angle was misaligned (averaged over all voxels in the ROI; Aligned > Misaligned, *t*(34) = 3.1, *p* = .004). To test the specificity of this effect, we also ran additional general linear models testing four control periodicities: four-fold, five-fold, seven-fold and eight-fold. For the five-fold and eight-fold test data, Shapiro-Wilk tests indicated that the data were not normally distributed (5-Fold: *W* = 0.865, *p* < 0.001; 8-Fold: *W* = 0.840, *p* < .001). We therefore used nonparametric Wilcoxon signed rank tests to determine whether the median contrast (grid-like coding) value differed from zero, Bonferroni-corrected for five tested periodicities (α = .01). Only six-fold symmetry differed significantly from zero (Wilcoxon *W* = 145, *p* = 0.005; Figure 2c). We further established the specificity of the six-fold effect by running a permutation test with 10,000 iterations to assess the likelihood that the overall pattern observed could have arisen by chance. For each participant, we randomly permuted their contrast values across the five periodicities, preserving the overall distribution but disrupting any effects. After each permutation, we recalculated Wilcoxon *W* for each periodicity using the permuted data, recording *W* and *p*-values for each periodicity. Then we checked whether a pattern as extreme as the observed six-fold significance (with permuted *W* at least as low as observed *W*) occurred while simultaneously all other periodicities remained non-significant. This pattern occurred in only 427 of the 10,000 permutations (*p* = 0.04).

**Figure 2:**
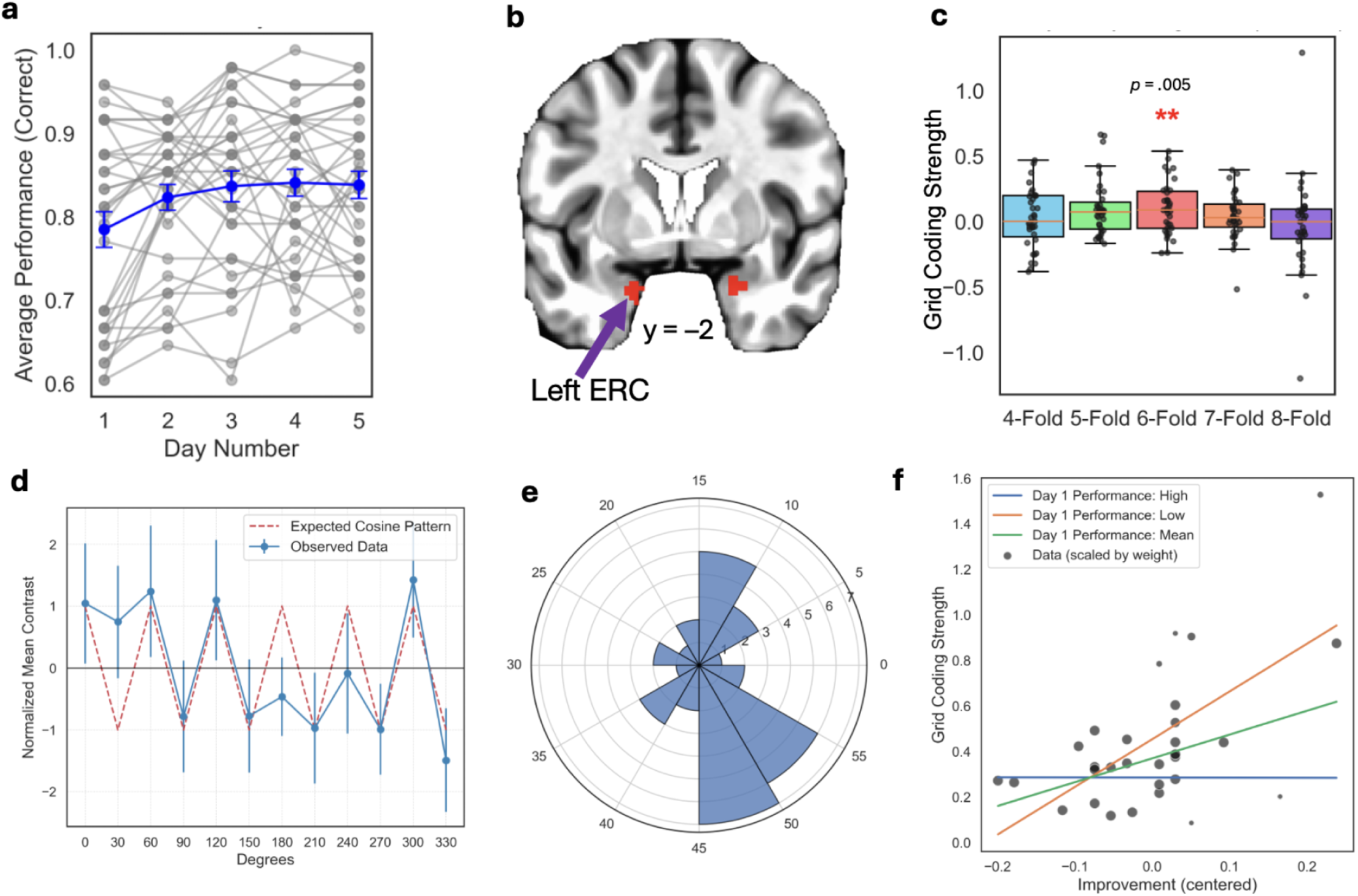
Grid-cell like activity pattern was detected in left entorhinal cortex during abstract auditory navigation. **(a)** Training task performance across the 5 days. (b) Results of GLM0 (six-fold symmetry localizer in entorhinal cortex) showing significant clusters (*p* < .005, corrected). **(c)** Contrast values (Aligned > Misaligned) for grid-cell-like activity in entorhinal cortex (six-fold) as well as control periodicities. **(d)** Activity (Aligned > Misaligned) by trajectory angle, aligned to participants’ preferred grid angle, revealing a sinusoidal pattern where contrast values rise for Aligned values (0, 60, 120, 180, and 300) and fall for Misaligned values. Bars indicate SEM. **(e)** Histogram plot of participants’ preferred grid angle φ in 60-degree space. **(f)** Grid-like coding strength was predicted by learning gains across the five training days. The effect was significant even after controlling for Day 1 performance and its interaction with Improvement. Green line indicates the slope for participants with average initial accuracy, orange line indicates the estimated slope with below average initial accuracy (-1SD), and blue for participants with high initial accuracy (+1SD). Point size is scaled to illustrate its weight in the robust regression model.

For BOLD signal to be consistent with grid-like coding, the signal should fall and rise in a sinusoidal pattern every 30 degrees, rising for the preferred grid angle and its 60 degree multiples, and falling for misaligned angles offset by 30 degrees. To test for this pattern, we separately plotted the mean contrast values in 30 degree increments. This revealed the predicted sinusoidal pattern (Figure 2d), the statistical significance of which was assessed by *z*-scoring each participant’s data and calculating a Pearson correlation with the expected pattern over the twelve angles from 0 to 300 degrees [𝑐𝑜𝑠(6θ)]. Correlation coefficients were converted to Fisher’s *z* and tested against zero with a one sample t-test, with results indicating a significant deviation from zero (*t*(34) = 4.09, *p* < .001). Analysis of participants’ preferred grid angles using a circular histogram revealed no evidence that the distribution was non-uniform (Rayleigh’s Test *Z* = 1.45, *p =* .23; Figure 2f). Having finished analysis of the left entorhinal ROI, we tested the right entorhinal ROI using the same procedure, but there was no evidence of grid-like coding aligned to a preferred grid angle across the even and odd runs in this region, using the procedure we had applied to left entorhinal cortex (Aligned > Misaligned, six-fold, *p* = .41). The right entorhinal cortex was therefore not analyzed further.

### Correlation with behavior

Having established robust grid-like coding in left entorhinal cortex, we next asked whether these signals predict individual differences in task performance, musical background, or subjective effort. The strength of grid-like coding was re-quantified by estimating a new model (GLM3) that was identical to GLM1 but included all functional runs, and estimating another model (GLM4), which was the same as GLM2, but with all runs included. The inclusion of all runs here was intended to improve the estimation of the preferred grid angle and neural activity during the task by leveraging the full dataset, given that the split-half analysis had already confirmed consistent six-fold symmetry. Because a Shapiro-Wilk test revealed that grid-like coding values deviated significantly from normality (*W* = 0.85, *p* < .001), we used a robust regression approach to mitigate the influence of outliers and non-normal errors. Specifically, we fit a robust linear model (RLM) with Huber’s T as the M-estimator, implemented via iteratively reweighted least squares (IRLS) in the *statsmodels* Python package. This procedure adjusts observation weights at each iteration, down-weighting extreme residuals to produce parameter estimates that are less sensitive to outliers compared to ordinary least squares (Huber, 1981; Wager et al., 2005). For the analysis, we created separate regressors for initial Day 1 performance (z-scored), Improvement (Day 5 minus Day 1, z-scored), and their interaction. This allowed us to test for an effect of improvement independently of the participant’s starting performance. We also included z-scored age, subjective online-training difficulty ratings, scanner-task difficulty ratings, and musical experience, calculated as the sum of the z-scores of years of musical training (i.e. private or group lessons), and years of musical experience (e.g. playing in a band or singing in a choir).

There were *N* = 30 participants with complete data. A significant negative interaction between Day 1 performance and Day 1 to Day 5 improvement emerged (*z* = -0.1, *z* = -2.9, *p* = .004; Figure 2f): participants starting with low performance exhibited a strong link between learning and grid coding, whereas participants already near ceiling on Day 1 showed a weaker relationship, as there was less room for improvement. However, despite this interaction, overall improvement also emerged as a significant main effect predictor on its own (β = 0.10, *z* = 2.3, *p* = .02), indicating that participants who improved more from Day 1 to Day 5 showed higher grid-coding strength, even when accounting for initial performance levels. Self-reported online training difficulty also significantly predicted grid-like coding (β = -0.07, *z* = -2.2, *p* = .03) such that higher perceived difficulty related to lower grid-coding strength— ruling out an alternative account whereby greater grid-like-coding was driven by increased effort. Other factors (scanner-task difficulty, age, and musical experience, initial performance) were not statistically significant. With 30 participants and 7 predictors, this analysis should be interpreted cautiously, and may not have detected small but real effects. A simpler model with only Day 1 performance and Improvement and their interaction was qualitatively similar (Improvement and the interaction both statistically significant).

We did not include final (Day 5) performance in the model due to multicollinearity (Day 5 performance being the sum of Day 1 performance and Improvement). To examine final performance as a standalone predictor, we fit a parallel model substituting performance on Day 5 (z-scored) for both baseline (Day 1) and improvement. Day 5 performance did not emerge as significant (*p =* .8), nor did any covariates. This outcome indicates that final performance does not meaningfully predict grid-coding strength—it is most strongly predicted by learning gains. Statistics for all predictors are reported in Table S1.

### Exploratory whole-brain analysis of grid-like coding signal

Our primary hypothesis concerned entorhinal cortex and we therefore restricted our analysis to this region. However, because previous studies of grid-like coding during spatial or conceptual navigation have identified additional regions exhibiting this signal, we report an analysis of GLM0 (six-fold ‘localizer’) within a whole brain gray matter union mask. For transparency and thoroughness of results, we report significant results at several thresholds: voxel-wise p < .05, .01, .005, .001, and .0005, each with a corresponding cluster extent threshold applied (FWE *p* < .05). Grid-like coding was detected in several regions outside of entorhinal cortex, most prominently the left temporoparietal junction, left dorsolateral prefrontal cortex, right dorsomedial and ventromedial prefrontal cortex, and right orbitofrontal cortex (Figure 3.)

**Figure 3:**
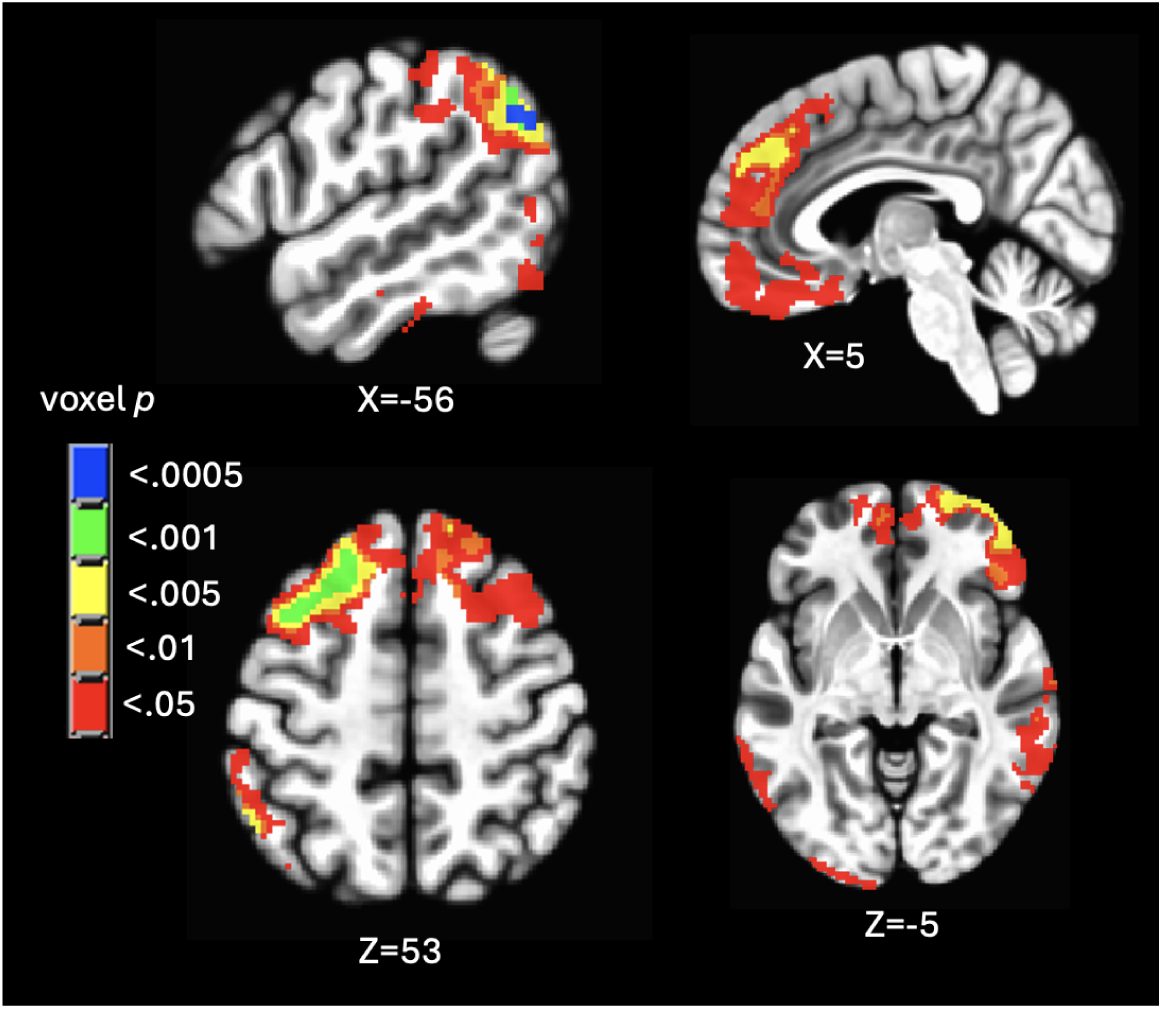
Exploratory whole-brain analysis of six-fold symmetry. Grid-like coding was detected in several regions outside of entorhinal cortex, most prominently left temporoparietal junction, left dorsolateral prefrontal cortex, right medial prefrontal cortex, and right orbitofrontal cortex. Results are presented across several voxel-wise thresholds, FWE cluster corrected at each threshold.

### Behavioral Interference (Experiment 2)

In Experiment 2, participants performed the auditory task with either a concurrent spatial or non-spatial interference task (Figure 4a). The spatial task required visually scanning the QWERTY keyboard to work out letter positions, while the non-spatial task required reasoning about serial alphabetical order. Auditory task responses were selectively slowed by the spatial interference during the Move condition (Movement X InterferenceType interaction F(1,40) = 4.61, *p* = 0.038, *η_p_*^2^ = .103; post-hoc *t*-tests at Bonferroni-corrected α = 0.025: Move_Spatial_ > Move_Non-Spatial_ *t*(40) = -2.6, *p* = 0.011; No-Move_Spatial_ > No-Move_Non-Spatial_ *t*(40) = -1.4, *p* = 0.157; Figure 4b). The magnitude of spatial interference was calculated as the difference in response times between the Spatial task and the Non-Spatial concurrent task, for the Move condition, and these values were plotted for each participant (Figure 4c).

**Figure 4:**
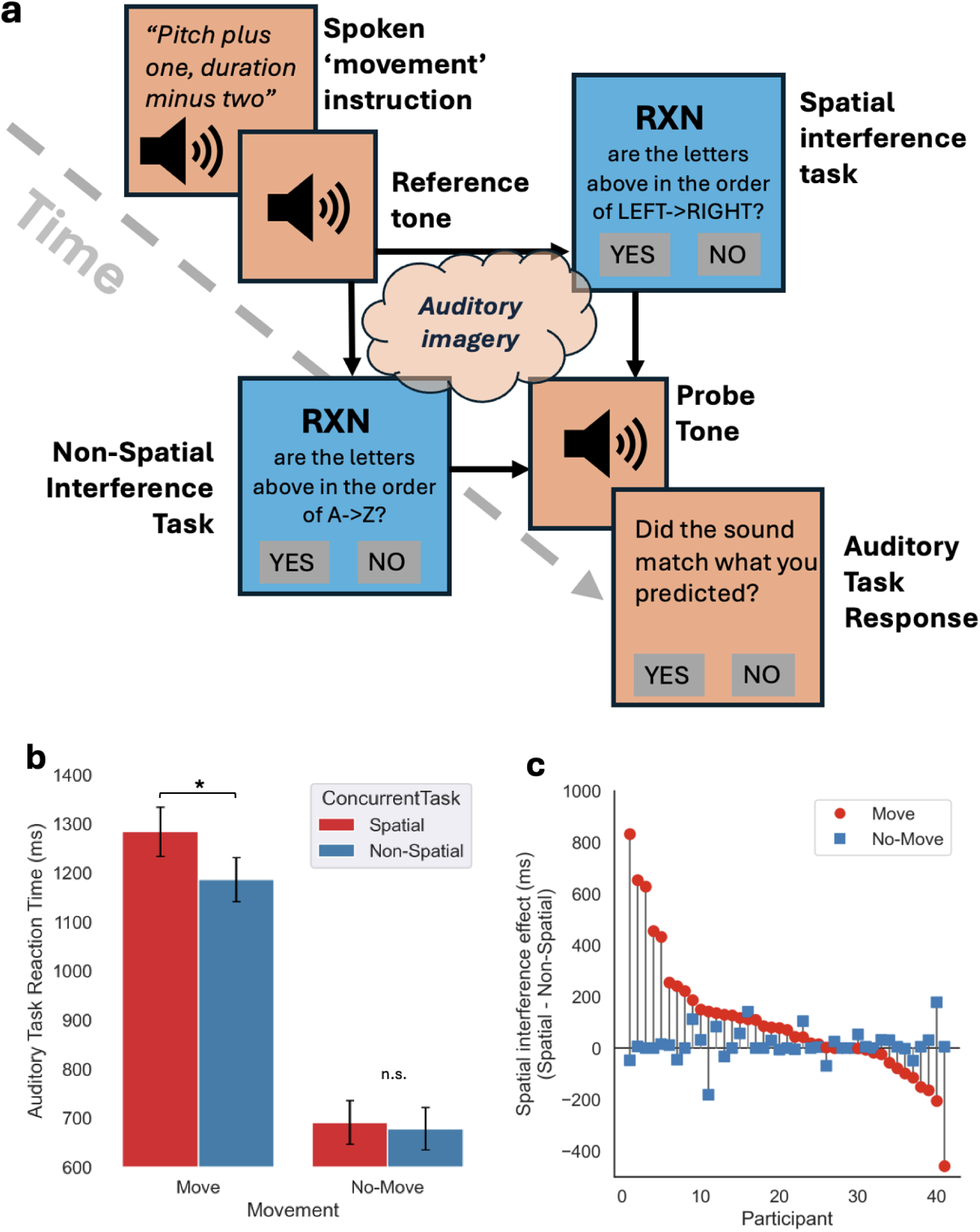
Behavioural experiment design. **(a)** The trial starts in the upper left, and continues to the bottom right. Peach boxes show main (auditory) task components and blue boxes show concurrent tasks. Trials began with a spoken transformation instruction, followed by the Reference Tone. Participants then actively imagined a new tone while simultaneously performing an interference task: they saw three capital letters and judged whether they were arranged left to right on the QWERTY keyboard (spatial condition), or whether the sequence was in alphabetical order (non-spatial condition).They then returned to the main task and immediately heard the probe and judged whether it matched their predicted (imagined) tone. **(b)** Response time for the auditory navigation task in the Move and No-Move condition, for both the Spatial and Control concurrent tasks. Participants’ judgments were slower when they needed to “move” in the auditory space (imagine a new tone) while simultaneously performing the spatial concurrent task. **(c)** Plot of the spatial interference effect for each participant, sorted by magnitude. The effect was defined as the difference in response times in the auditory task (Spatial concurrent task – Control concurrent task). Red = Move, Blue = No-Move.

Although performance accuracy was not a measure of interest, we report that participants were overall more accurate for the No-Move trials (97%) than the Move trials (78%; F(1,40) = 92.3, *p* < .001), but there was no significant difference in accuracy as a function of Interference Type, nor an interaction of Movement and Interference Type (Figure S1 1; Table S2).

We also analyzed performance on interference tasks themselves: accuracy was significantly higher in the Spatial condition (97%) than the Non-Spatial condition (94%; F(1,40) = 5.14, *p* < .029). This suggests the Spatial task was slightly easier than the Non-Spatial task, ruling out the possibility that the effect of Spatial interference could be explained by increased difficulty. Interference task response times overall were slower when the auditory task was in the Move condition (F(1,40) = 61.4, *p* < .001). There were no other significant effects (Table S1).

## Discussion

In two experiments, we found convergent evidence that mental navigation in a purely auditory pitch-duration space recruits core spatial coding mechanisms in the entorhinal-hippocampal system. Experiment 1 used fMRI to reveal robust six-fold grid-like modulation in left entorhinal cortex when participants imagined transforming tones within a pitch-duration space. Notably, participants with stronger grid-like activity showed greater learning gains over five days of training, suggesting that spatial coding mechanisms facilitate the acquisition of abstract auditory knowledge. Experiment 2 used a behavioral interference paradigm to test whether these spatial resources are causally necessary for task performance. Performance was selectively disrupted by a spatial interference task only when participants had to mentally navigate (i.e., move) in the auditory space, indicating that spatial neurocognitive resources underlie these conceptual transformations.

These findings extend prior work on grid-like representations in humans, which has demonstrated entorhinal six-fold symmetry during navigation in physical space and non-spatial domains—primarily visual, olfactory, or social (e.g., Constantinescu et al., 2016; Bao et al., 2019; Park et al., 2021; (Sigismondi et al., 2024). These findings collectively support the notion that entorhinal coding may be domain-general yet shaped by experience. By contrast, our results provide a clear demonstration of grid-like coding in a purely auditory domain and, crucially, show that interfering with spatial processing impairs auditory conceptual navigation. Thus, the present study offers a novel advance: it links the neuronal code observed in entorhinal cortex for abstract tasks with causal behavioral evidence that these spatial mechanisms are essential to mentally transform and learn information even in non-visual, multidimensional auditory contexts.

The correlation between grid-like coding in left entorhinal cortex and training-related improvement highlights a potentially broader role for spatial coding mechanisms in learning. Grid cells may optimize learning by providing a structured coordinate system for newly acquired information, whether that information corresponds to locations in a physical maze or tones in a conceptual “pitch–duration” space. Participants who show stronger grid-like signal with fMRI may have a stronger and more consistent representation to scaffold learning, though an alternative is that both enhanced entorhinal signal and better learning stem from a common underlying cause. Longer-term, repeated measurements would help clarify this relationship. Moreover, our findings align with domain-general accounts positing that place- and grid-like responses can emerge from a general clustering or concept-learning process (Mok and Love 2019). In such a framework, grid-like patterns may arise when uniformly sampling a low-dimensional space, reflecting learning dynamics rather than a strictly spatial representation.

Future work might also include investigating whether such grid-like signals emerge in more naturalistic auditory or auditory-motor tasks, such as musical production where multiple acoustic dimensions (pitch, loudness, timbre) must be navigated simultaneously, and whether the magnitude of grid coding predicts individual differences in skill or expertise. In principle, exploring auditory spaces with three or more dimensions (e.g., pitch, duration, and timbre) could further reveal how grid coding scales with complexity. Learning of new linguistic features, such as speech contrasts conveyed acoustically across multiple dimensions (i.e. fundamental frequency and duration) could also rely on similar coding mechanisms, and could be an area of further exploration.

An important caveat is that Experiment 1 was not designed to establish a direct causal link between entorhinal grid-like activity and auditory conceptual transformations; the correlation with learning outcomes is strong but does not by itself demonstrate entorhinal cortex necessity. Our spatial interference study (Experiment 2) supports the conclusion that spatial resources are required, yet cannot localize the disruption explicitly to entorhinal cortex. Causality could be demonstrated more strongly in the future using other methods, such as intracranial stimulation.

Our findings demonstrate that spatial neurocognitive resources causally support auditory conceptual navigation, consistent with a role for the hippocampal-entorhinal system in sound processing (Billig et al., 2022). Grid-like coding in the left entorhinal cortex predicts learning, suggesting this ancient navigation system facilitates the acquisition and manipulation of multidimensional knowledge. These results highlight the versatility of spatial coding in encoding diverse forms of information and suggest broad applications for understanding learning and expertise across complex, real-world tasks.

### Methods Experiment 1 (Behavioral training and fMRI)

#### Participants

Thirty-five participants (M=23, F=12) completed the study in exchange for payment of GBP £50. The experiment consisted of 5 days of at-home training, followed by an in-person scanning session at the Birkbeck-UCL Centre for Neuro-Imaging. Participants gave informed consent in a protocol approved by the UCL Psychology Ethics Steering Committee. The majority of participants were recruited only from the UCL and Birkbeck student and staff community, in line with UCL’s COVID-19 safety guidelines that were in place during testing in 2021.

#### Stimuli

The stimulus space consisted of 36 complex tones that varied in duration and frequencies (Figure 1a). The sounds were based on “Risset’s bell” which is a synthesized bell-like sound composed of 11 sinusoids (Risset & Wessel, 1999). The ratios of the frequencies of sinusoid components are designed to be non-harmonic so as not to give rise to a single pitch percept, thereby deterring music-based task strategies. The tones were 40ms, 80ms, 160ms, 320ms, 640ms, or 1280ms in duration, with 10ms cosine onset and offset ramps to avoid transients. The frequency profile of the bells was adjusted by shifting the global frequency (G) and generating sinusoids that were multiples of this value. The G values were 260Hz, 343Hz, 453Hz, 597Hz, 788Hz, and 1040Hz (Figure 1b). Sinusoids making up the bell sounds were calculated as in Supplemental Table 1. Unlike the classic Risset bell, the sinusoid components did not decay, instead maintaining constant amplitude over time until the off ramp.

A set of 48 ‘trajectories’ was created that effectively sampled all angles through the pitch–duration space at approximately 15 degree increments (Figure 1c). A trajectory was composed of a starting tone (the “reference tone”) and a final tone at the end of the trajectory (the “probe tone’). The probe tone correctly reflected the transformation instructions (Match trials). Mismatch trials were created by semi-randomly modifying the probe tone to closely align with the intended trajectory instructions (within the same quadrant as the matched trajectory) without fully matching them, ensuring participants perceived these as similar to the imagined target but not identical (Mismatch Trials).

#### At-Home Training

The training task was created and deployed using Gorilla Experiment Builder (Anwyl-Irvine et al., 2020). Participants underwent five consecutive days of training in order to allow thorough learning of the auditory space and achieve the highest performance possible ahead of the fMRI scanning session, which naturally would bring more challenges and distraction due to awkward physical positioning and scanner noise. Extensive training also ensured consistent mental representation, increasing the chances we could detect stable grid-like coding. On the first day, participants listened to a spoken audio recording that introduced the participant to the space and explained the instructions. The instructions were carefully designed not to include any spatial language (such as *high* or *low*) that could lead participants to adopt an explicitly spatial strategy.

The task was designed to be analogous to previous tasks that have examined grid-like coding of conceptual navigation (Bao et al., 2019; Constantinescu et al., 2016). Participants first heard a reference tone, followed by verbal instructions indicating a trajectory: The word “pitch” or “duration” was spoken, followed by *“minus [x]” or “plus [x]”* where *x* was the number of levels by which the participant would need to mentally adjust the starting tone to achieve the ending tone, or the word “same” to indicate that change was not required for that particular dimension. For instance, *“Pitch, minus one. Duration, plus two”* indicated to the participant that they would need to imagine a new sound that is one rank lower in pitch, and two ranks longer in duration. *“Pitch, same. Duration minus one”* indicated that the new sound to be imagined should have the same pitch but be one rank shorter in duration. The purpose of using numbers was to avoid explicitly spatial words such as higher, lower, longer or shorter. Participants then mentally imagined a new tone based on the reference tone and trajectory. After they had done this at their own pace, they pressed the spacebar. Next, they heard the probe tone. If the probe tone matched the sound they had imagined, they indicated this by pressing the “K” key. Otherwise, they pressed the “J” key. On each day of training, participants first completed 24 trials where all trials were Matches, to familiarize themselves with the tone space. Following this they completed 48 trials for which only 50% of the trials were Matches (the rest Mismatches). Scores were not logged for four participants due to software errors.

#### In-scanner

On Day 6, the task used in the scanner was similar to the training task except 1) it was not self-paced; rather, the participant imagined the tone during a four-second interval following the reference tone, and 2) participants were informed that the vast majority of trials (∼88%) were ‘matches’ and they should stay vigilant and only press a key (with index finger on a keypad) for a rarely-occurring ‘mismatch’ trial. The reason for this choice was to minimize response-related motion and encourage participants to simply focus on imaging the target tone without worrying about performance. Responses to the minority vigilance trials were not logged due to a technical issue, but keypresses were visually confirmed and alertness and engagement monitored throughout, using an in-scanner camera. Participants completed four runs of 48 trials (9.58 minutes per run). Each trial began with the spoken trajectory instruction and a starting tone, followed by a 4-second “imagery” period during which participants imagined what the probe tone should sound like, followed by presentation of the probe tone, then 2 – 2.2 second period before the next trial. A minority of participants (*n* = 4) completed only three runs. After scanning participants completed a questionnaire about their experience with the task and musical background.

#### MRI Data Acquisition

Participants were scanned with a Siemens Prisma 3.0 Tesla MRI scanner using a 32-channel head coil. Functional images were acquired using a T2*-weighted echo planar image pulse sequence (44 oblique axial slices, in-plane resolution 3mm^2, 3mm slice thickness, no gap, TR = 1250 ms, TE = 41 ms, flip angle = 61°, matrix size = 64x64, FOV =192 mm). All EPI functional scans were performed using 4 multiband acceleration (Feinberg et al., 2010; Feinberg & Setsompop, 2013). There were 460 volume images acquired per run, with the first 8 images discarded to allow for longitudinal magnetization to arrive at equilibrium. Sounds were presented using Sensimetrics S14 earbuds, padded around the ears with NoMoCo memory foam cushions. High-resolution T1-weighted structural images were acquired using an MPRAGE sequence with isotropic 1 mm voxels in a 256 × 256 × 176 matrix.

#### MRI Pre-Processing

Functional MRI data were preprocessed using SPM12 (Ashburner et al., 2014). The first eight volumes of each run were discarded to eliminate the pre-steady state period, ensuring the stability of the fMRI signal. The remaining images were realigned to the mean image of each run to correct for head motion. Following realignment, the functional images were co-registered to the corresponding participant’s anatomical T1-weighted image. The anatomical images were then segmented into gray matter, white matter, and cerebrospinal fluid (CSF) using SPM12’s segmentation algorithm. The resulting forward deformation field from the segmentation process was applied to warp both the anatomical and functional images into standard Montreal Neurological Institute (MNI) space, reslicing them to 2mm isotropic voxels. Finally, the functional images underwent spatial smoothing with a Gaussian kernel of 6mm full-width at half-maximum (FWHM) to improve the signal-to-noise ratio and to account for anatomical variability among participants. Overall motion was assessed by calculating average frame displacement (FD)—the sum of the absolute temporal derivatives of the six motion parameters, with rotations converted to distances using an arc length on a 50 mm radius sphere (Power et al., 2012). Motion was on average low (mean FD < 0.19 ± 0.06 mm). It is highly implausible that small, random head movements could spuriously generate a systematic 60° modulation of the BOLD signal aligned with trial-by-trial pitch-duration angles, mimicking or inflating grid-like patterns specifically in our region of interest. Therefore, given that excluding any runs could compromise our split-half design and statistical power more than it would reduce potential motion effects, we opted to retain all the data for analysis.

#### Mask and ROI definition

Our central hypothesis was that six-fold grid-like coding would occur in entorhinal cortex. We therefore restricted our primary analysis by creating a mask of voxels in this region bilaterally. The mask was created using the *aparc+aseg* Freesurfer parcellations (Fischl, 2012) for left and right entorhinal cortex for each participant, which were projected into MNI space using each participant’s forward deformation field obtained during preprocessing. The participant-level masks were then summed and binarized to create the group union mask. For an exploratory whole-brain brain analysis we report at the end of the results, a gray matter mask was defined as the union of individual participant gray matter masks from SPM preprocessing, in MNI standard space.

#### Six-fold localizer within entorhinal cortex: GLM0

Tests for six-fold grid-like coding was performed using a standard procedure first developed by Doeller et al., 2010, implemented with the GridCAT Tool box (Stangl et al., 2017) in MATLAB (Matlab, 2012), wherein regions with potential grid-like coding are first localised with a quadrature filter, then, in an orthogonal analysis, all data is split into a Training Set and Test Set for independent validation.

First, a quadrature filter was deployed to localize voxels within entorhinal cortex that showed the strongest six-fold signal. It included regressors for the stimulation presentation period, as well as the ‘mental navigation’ period. Two additional regressors were added to the GLM model calculated as cos6θ and sin6θ, where θ is the angle of the transformation trajectory for a given trial (Constantinescu et al., 2016; Doeller et al., 2010). This method decomposes the trajectory angle of each trial into *x-* and *y-*axis components on a unit circle, with six-fold periodicity, capturing the six-fold modulation characteristic of grid-cell activity. Because T2* signal in the medial temporal lobe is notoriously weak due to high susceptibility, the implicit masking threshold was switched off (-Inf), allowing for the inclusion of all voxels in the analysis regardless of their initial intensity values (Nau, 2019). High-pass filtering was implemented with a cutoff of 1/180 Hz for each run to remove low-frequency drifts from the data. The spatial smoothness of the residuals of these models was measured with AFNI’s *3dFWHMx* for follow-up cluster extent correction using the ACF model.

From the cos6θ and sin6θ regressors we obtained parameter estimates βcos and βsin. Rather than assess six-fold symmetry *F*-tests on the subjects’ parameter estimates, which then need to be converted to *Z* at the participant-level for group analysis (Bao et al., 2019; Constantinescu et al., 2016), we instead included both coefficients together for each run and each participant in a linear mixed effects model (using AFNI’s *3dLMEr* command) with Beta (βcos and βsin) as a fixed effect and Participant as random intercept, effectively modelling all the data in a single step. An *F*-test was then conducted on the joint effect of the coefficients (1*βcos & 1*βsin), and results were thresholded voxelwise at *p* < .001, cluster corrected to *FWE p* < .05 using AFNI’s *3dClustSim* (10,000 iterations, with ACF model; minimum cluster size 10 voxels). This analysis localized voxels that were sensitive to either of the cos6θ and sin6θ regressors, indicating six-fold rotational symmetry. Crucially, this analysis identifies six-fold symmetry in voxel responses regardless of grid angle orientation, and is therefore orthogonal to GLM1 and GLM2 below, which estimate and test specific grid angles on a participant-wise basis.

#### Grid angle estimation (GLM1) and validation (GLM2)

After identifying candidate voxels with a quadrature filter, we then split the data into Training and Test sets to estimate and validate the preferred grid orientation in an independent manner. In line with previous studies, we used a split-half approach (Constantinescu et al., 2016; Doeller et al., 2010), where grid angle sensitivity is estimated in one partition of the data (Training Set), and tested in an independent partition (Test Set). This allowed us to detect the presence of grid-like coding while avoiding bias. Two additional GLMs were created. GLM1 included parametric sine and cosine regressors for only odd-numbered runs for each participant, and was used to estimate βcos and βsin coefficients as above. The coefficients were then used to calculate the ‘preferred’ grid angle φ across the odd runs:

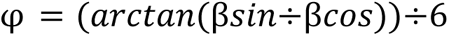

GLM2 tested for 60° sinusoidal modulation of the BOLD signal in the Test Set (even runs) that was *specifically aligned* with the preferred grid orientation identified in the Training Set (odd runs). This was done by calculating the cosine of the angle of each trial’s trajectory θ after being realigned with the participant’s preferred grid angle φ:

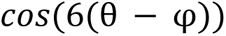

Trials were then coded as ‘Aligned’— defined as the trajectory angle lying within ±15 degrees of the preferred grid orientation or a 60 degree multiple thereof—or ‘Misaligned’, referring to all other angles (Stangl et al., 2017)(Figure 1d, 1e). Then we tested for greater activation for Aligned than Misaligned trials within the independent Test Set (GLM2).

### Experiment 2: Spatial processing interferes with a navigating a conceptual pitch–duration ‘space’

#### Participants

Fifty-one participants (with no overlap from Experiment 1) were recruited for the study, 16 internally at Birkbeck College and 35 through Prolific, and paid £5 for 30 minutes of participation. Data from ten participants were excluded for not meeting an *a priori* performance threshold of 60% correct responses on either the main or interference task (corresponding to a lenient *p* < .10 relative to 50% chance level on each task). As performance and engagement of online participants can be highly variable, this threshold ensured that we only included participants who demonstrated the ability to perform the tasks beyond mere guessing (Jasmin et al., 2021; Support, 2020). The included participants were aged 18 – 45 (M = 28, SD = 7.3), 10 male, 30 female, and 1 identifying as non-binary. All participants were fluent in English, educated to at least UK A-level standard, and had no known impairments to their listening, reading, and writing capabilities such as dyslexia, blindness, and deafness.

#### Stimuli

Six ‘trajectories’— combinations of starting tones and transformation instructions—were selected from Experiment 1 to sample the auditory stimulus space. These trajectories appeared with both a Matching probe tone and Mismatching probe tone during the experiment. These constituted the “Move” condition of the experiment. Next, “No-Move” versions of these trials were created by taking the 12 Move trials, keeping the reference tone the same, and replacing the transformation instructions with new instructions that required the participant to imagine the same tone again. Participants therefore only needed to maintain in auditory working memory the reference tone, and judge whether the probe tone matched it. Each of these 24 trials was presented twice in the experiment, once with the Spatial interference task and once with the Non-Spatial interference task. This produced 48 total trials that fully counterbalanced the Match vs Mismatch and Move vs No-Move conditions as well as the Spatial vs Non-Spatial interference tasks.

#### Main Task

As in Experiment 1’s training task, participants were first presented with instructions that familiarized them extensively with the stimulus space, exposing them to the range of sounds they would hear and the various levels of the pitch and duration dimensions. Instructions were followed by a short practice session that gave the participant feedback on their responses. In the main experiment, participants were first presented with a fixation cross before hearing spoken instructions on how the upcoming tone should be manipulated. As described above, there were two types of trials: Move and No-Move. The Move trials required mental ‘navigation’ through the stimulus space, while the No-Move trials merely required maintaining the first sound in memory. On Move trials, participants heard a male voice speak a prompt that took the following form: The word “pitch” or “duration” was spoken, followed by *“minus [x]” or “plus [x]”* where *x* was the number of levels by which the participant would need to adjust the starting tone to achieve the ending tone, or the word “same” to indicate that change was not required for that particular dimension. In trials without movement (No-Move), participants always heard “*Pitch, same. Duration, same*”, prompting them to simply imagine the same sound they had heard.

The verbal instructions were immediately followed by the presentation of the ‘reference tone’, giving the participant a starting ‘location’ within the auditory space. As the participants began to mentally imagine the target sound based on the starting sound and transformation instructions, the interference task began (Figure 5a; described below). After responding to the interference task, the participant immediately heard the final sound (the ‘probe tone’) which either matched or mismatched the target sound they were instructed to imagine. The participant responded using the J (match) and K (mismatch) keys on the keyboard, then received auditory “correct” or “incorrect” feedback on their response. The order of trials was randomized.

#### Interference Tasks

There were two interference tasks—one spatial and one non-spatial—that used the same stimuli. Because our auditory space was two-dimensional, for our spatial interference task we used another two-dimensional space, namely, the QWERTY keyboard layout. Previous work in humans has shown using fMRI that scanning visual space recruits a six-fold grid-like coding within entorhinal cortex (Julian et al., 2018; Nau et al., 2018). We posited that scanning up, down, left and right across the 2D QWERTY keyboard to work out relative letter positions should also recruit grid-like neural codes and spatial neural resources, and only interfere with the 2D conceptual auditory space if it is processed with overlapping resources.

Each stimulus consisted of three capital letters (that did not form words), presented visually, to which the participant responded *YES* or *NO* with a keypress. In the Spatial task, participants judged whether or not the letters shown were in LEFT → RIGHT order in the QWERTY keyboard layout, requiring 2D visual scanning. In a Non-Spatial control task, participants were asked to simply judge whether the letters shown were in alphabetical order (A → Z) in the English alphabet. While some cultural artifacts do lay out the alphabet spatially, the extent to which individuals show spatial associations with alphabetical order is limited and idiosyncratic (Jonas et al., 2011). Moreover, previous neural data during alphabetic order judgments have not implicated the hippocampal-entorhinal system (instead showing anterior intraparietal sulcus activity) (Attout et al., 2014). All materials are openly available on Gorilla Open Materials. The experiment was carried out using the Gorilla platform (Anwyl-Irvine et al., 2020) and was approved by the Psychology Department Ethics Committee at Birkbeck, University of London.

#### Analysis

The primary measure of interest was cognitive load as assessed by response times on the main task. The interference tasks were designed to be easy enough that incorrect responses could be assumed to be due to inattention rather than inability. Therefore we only analyzed median response times for each participant for which the interference task responses were correct—the vast majority of trials (∼96%). Two-by-two repeated measures ANOVAs were performed using the response times as the dependent variable, with independent variables Movement (Move vs. No-Move), Interference (Spatial vs. Non-Spatial), and their interaction. An analogous analysis was also performed with accuracy scores.

## Data Availability

Anonymized data supporting the conclusions of the study will be made available in Birkbeck Institutional Research Online upon publication. The experimental paradigms are published on Gorilla Open Materials (Experiment 1: https://app.gorilla.sc/openmaterials/979793; Experiment 2: https://app.gorilla.sc/openmaterials/980195)

## Acknowledgments

Staff on this project were supported by an Early Career Fellowship from the Leverhulme Trust awarded to K.J. (grant number ECF-2017-151), and by an Arnold Bentley New Initiatives Fund award from the Society for Education and Music Psychology Research (SEMPRE) to K.J. We gratefully acknowledge the Birkbeck-UCL Centre for Neuroimaging (BUCNI) for providing MRI facility access. We also thank Andrew Persichetti for helpful comments on an early draft.

