## Supplement for "Spatial coding supports auditory conceptual navigation"

### Table S1: Descriptive statistics of regression variables

| Variable | Mean (SD) | Range |
|---|---|---|
| **Age** | 31.333 (7.294) | 20 – 52 |
| **Grid-Like Coding Strength** | 0.445 (0.307) | -0.008 – 1.548 |
| **Day 1 Performance** | 0.789 (0.118) | 0.604 – 0.958 |
| **Day 5 Performance** | 0.841 (0.092) | 0.667 – 0.979 |
| **Improvement (Day 5-Day1)** | 0.053 (0.098) | -0.146 – 0.292 |
| **Musical Training (Years)** | 8.000 (8.004) | 0.000 – 25.000 |
| **Musical Experience (Years)** | 9.333 (10.367) | 0.000 – 37.000 |
| **Online Training Difficulty Rating** | 2.667 (1.322) | 1.000 – 5.000 |
| **Scanner Difficulty Rating** | 4.467 (1.432) | 1.000 – 6.000 |

### Table S2: Experiment 1 Inferential Statistics

| Dependent Variable | Effect | Statistic | df | p Value | Significance |
|---|---|---|---|---|---|
| Main Task Reaction Time | Movement | F = 39.31 | (1, 40) | < .001 | *** |
| | Interference Type | F = 9.24 | (1, 40) | 0.004 | ** |
| | Movement by Interference | F = 4.61 | (1, 40) | 0.038 | * |
| Main Task Accuracy | Movement | F = 92.31 | (1, 40) | < .001 | *** |
| | Interference Type | F = 1.32 | (1, 40) | 0.258 | ns |

| | | | | | |
|---|---|---|---|---|---|
| | Movement by Interference | F = 0.13 | (1, 40) | 0.722 | ns |
| Interference Task Reaction Time | Movement | F = 61.44 | (1, 40) | < .001 | *** |
| | Interference Type | F = 2.84 | (1, 40) | 0.1 | ns |
| | Movement by Interference | F = 0.17 | (1, 40) | 0.679 | ns |
| Interference Task Accuracy | Movement | F = 0.07 | (1, 40) | 0.8 | ns |
| | Interference Type | F = 5.14 | (1, 40) | 0.029 | * |
| | Movement by Interference | F = 0.00 | (1, 40) | 1 | ns |

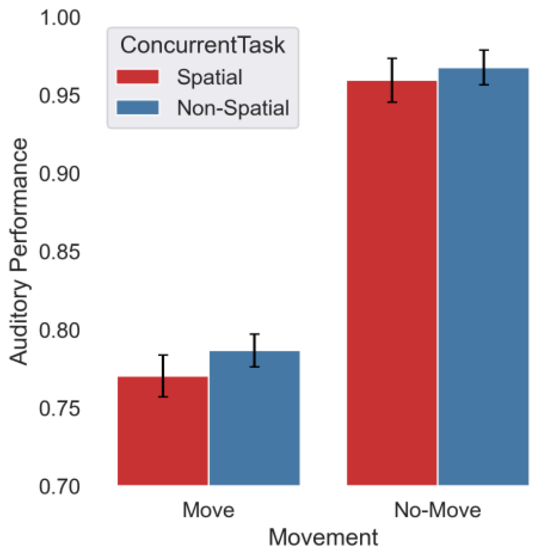


**Figure S1: Accuracy on Experiment 1.** Accuracy was higher for the No-Move condition, but there was no statistical difference between Spatial and non-Spatial concurrent task.
